# Family identity is represented within vocal categories by the gerbil auditory cortex

**DOI:** 10.1101/2025.10.24.684438

**Authors:** Estelle E. in ‘t Zandt, Ralph E. Peterson, Dan H. Sanes

## Abstract

Vocalizations are integral to social communication, with acoustic features that may convey both meaning and information about an animal’s identity or emotional state. The neural representation of vocalizations must therefore permit animals to generalize across one set of acoustic features to recognize meaning, and a different set of features to recognize vocalizer identity. To test this idea, we recorded the responses of gerbil core auditory cortex (AC) neurons to a large array (*n*>1500) of variants drawn from four vocalization categories and produced by 5 different families. Each vocalization category could be decoded from AC population activity with high accuracy, despite the acoustic variance across vocalization renditions or different families-of-origin, and displayed a ‘boundary effect’ (i.e., better classification across, than within, a category). Moreover, the family identity of each vocalization category could be decoded from a larger AC population. Thus, AC activity can be used to simultaneously predict vocalization category and family identity, and is robust to the natural range of acoustic variance.

## INTRODUCTION

Vocalizations often convey many pieces of information simultaneously. For example, spoken words convey meaning, yet also provide pivotal information about an individual’s identity (Abercrombie, 1967; Locke, 2021). In fact, it is acoustic variance that allows any single speech sound to convey both meaning and social information (Kleinschmidt, 2019; Kreiman et al., 2015; Lavan et al., 2019; J. J. Lee & Perrachione, 2022; Y. Lee et al., 2019). If the only acoustic variance was along an axis conveying category, categorization would be trivial while identification would be impossible. Variance thus increases the informational capacity of speech, yet creates a challenge for auditory processing: the nervous system must represent meaning associated with a vocal category by generalizing across interspeaker acoustic variance, while also representing social identity by discriminating between individual variance. Thus, to understand how the nervous system represents both vocalization categories and identity information, it is essential to use a large array of vocalizations with a natural range of acoustic variance (Herrnstein, 1990; Lavan et al., 2019; Louder et al., 2019).

Many animal vocalizations can convey both meaning and social identity, relying on aural communication to navigate complex social landscapes. For example, alarm calls signal the presence of nearby danger and ultrasonic calls signal emotional state (Knutson et al., 2002; Seyfarth et al., 1980). Acoustic variance also provides identifying information about individuals, including group membership (Barker et al., 2021; DeCasper & Fifer, 1980; Elie & Theunissen, 2018; Miller, 1979; Szenczi et al., 2016). The acoustic cues that convey this information are often multiple features (e.g. frequency, frequency modulations), and may differ across vocal categories (Elie & Theunissen, 2018; Kar et al., 2022; Kreiman et al., 2015). Thus, natural vocal processing is a complex operation that involves simultaneous extraction of both meaning and identity information from a variable set of acoustic signals.

Robust responses to vocalizations are measured throughout the auditory neuraxis (Carruthers et al., 2015; Gaucher et al., 2013; Lawlor et al., 2025; Meliza & Margoliash, 2012; Robotka et al., 2023; Russ et al., 2007; Schneider & Woolley, 2013), yet highly selective single neurons are uncommon (Glass & Wollberg, 1983; Wang & Kadia, 2001). Rather, the representation of vocalization categories is best explained by population codes (Carruthers et al., 2015; Montes-Lourido et al., 2021a; Robotka et al., 2023; Wang et al., 1995). For example, an auditory cortex (AC) population code can independently discriminate phonemes (Mesgarani et al., 2008) and social traits, such as age, from vocalizations (Shepard et al., 2015) in the presence of stimulus variance. While these studies point to AC as a vocalization processor, they characterize the processing of either vocal category *or* social information, but not together. In contrast, naturalistic social communication requires the simultaneous processing of both meaning and identity information. To date, it is unknown whether and how the AC is able to represent social information across many categories, and if so, what properties of the population code might underlie this representation.

Mongolian gerbils (*Meriones unguiculatus*) are a highly social species, with a rich repertoire of acoustically-distinct vocalizations that are associated with unique behaviors (Kobayasi & Riquimaroux, 2012; Ter-Mikaelian et al., 2012). Further, it has been previously shown that the usage of this repertoire differs between families (Peterson et al., 2024), posing the gerbil as an ideal candidate to study the neural mechanisms underlying simultaneous extraction of both meaning and social identity. Here, we ask whether gerbil AC neuron populations can simultaneously support vocal category discrimination, while also retaining the ability to distinguish between the family-of-origin for each vocalization. To test this, we performed wireless recordings from populations of AC single neurons in adult Mongolian gerbils while presenting >1500 vocalizations drawn from four different categories and 3 different families.

## RESULTS

### Gerbil vocalizations can be acoustically classified by category and family identity

To test whether auditory cortex (AC) activity can be used to categorize gerbil vocalizations, we required a large stimulus set that captured a comprehensive range of natural acoustic variance. Since gerbils exhibit family-specific usage of vocalization categories (Peterson et al., 2024), we obtained our stimulus set by recording vocalizations from 5 separate gerbil families (2 parents, 7-9 pups, postnatal (P) day 25-27). Vocalizations were first identified using a supervised deep neural network (DAS; (Steinfath et al., 2021) trained to identify individual, non-overlapping vocalizations (model performance F1 score: 0.77; **Fig S1A-B**). An unsupervised approach was then used to quantitatively characterize all vocalizations, allowing us to cluster them and ask in an unbiased way whether their acoustic features differed across families. Spectrograms of each individual syllable (i.e. variants) were embedded in a 32-dimensional latent space using a variational autoencoder (**Fig S1C-D**; see STAR METHOD; (Goffinet et al., 2021). A Gaussian mixture model was applied to these latents to generate clusters with similar acoustic attributes. A subset of clusters were then manually merged into broader groups that replicated vocal categories described in Peterson et al. (2024). Our experiments focused on 4 vocalization categories: combination (Combo), downward frequency modulated (DFM), upward frequency modulated (UFM), and Warbles recorded from 5 gerbil families (**Fig 1A-B**). We extracted a total of 50-250 variants per family per category, yielding a stimulus set of >2500 unique variants across the 5 subject animals (of which ∼1500 were used in any single recording session).

**Figure 1:**
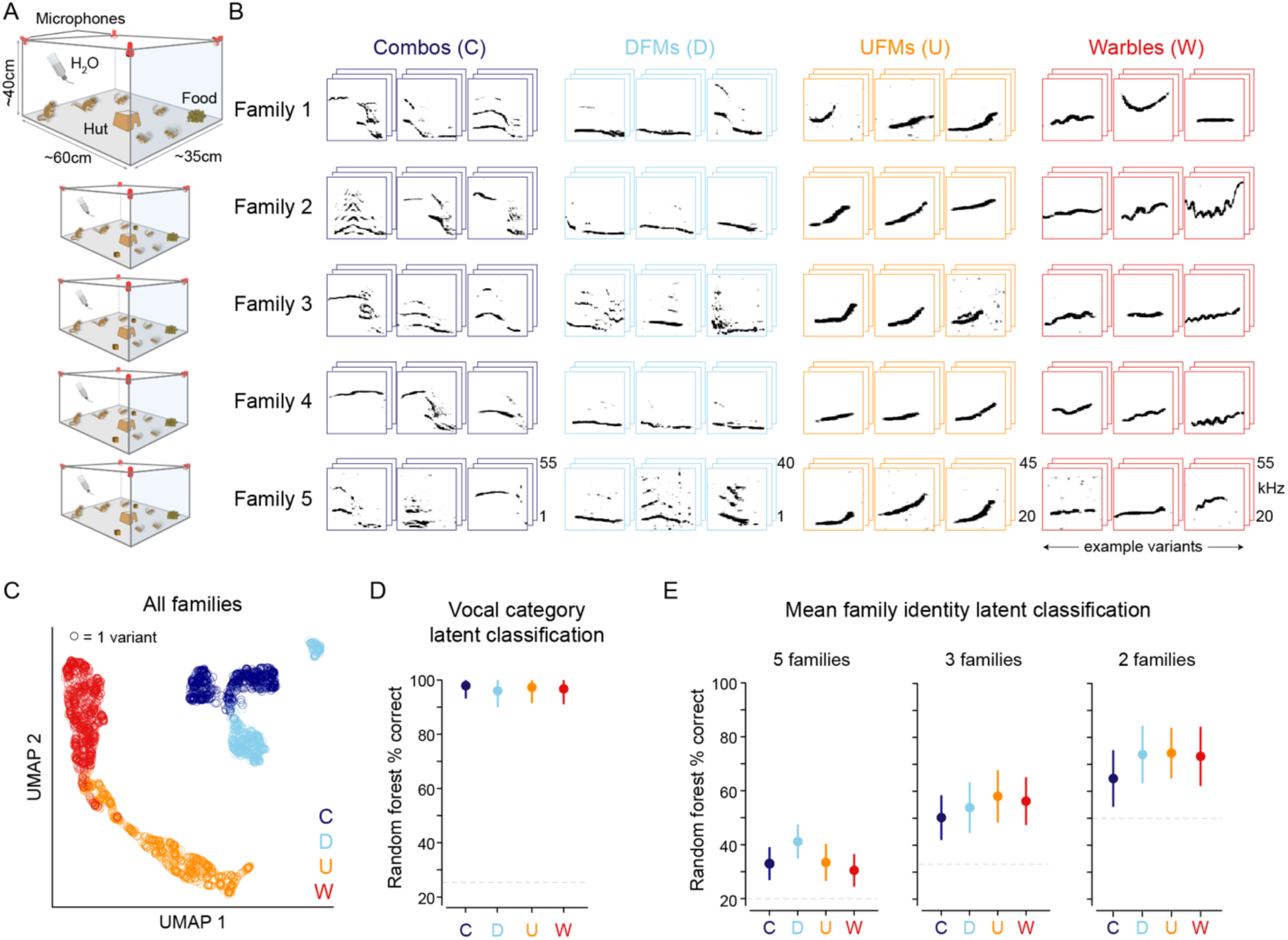
The acoustic attributes of gerbil vocalizations can be classified by both category and family identity. **A.** Schematic of enlarged home cage used for overnight vocalization recordings. **B.** Spectrograms of variants from each of 4 vocalization categories from each of 5 families. **C.** UMAP representation of vocal latents. Colors correspond to class labels in B. **D.** Random forest classifier performance on predicting vocalization categories trained on the vocalization latents. Error bars = 1 standard deviation (SD) across 100 SVM iterations. **E.** Random forest performance on predicting family identity trained on the vocalization latents. Classifier performance shown for 5, 3, and 2 families’ vocalizations. Family-ID classifiers were trained on a single category at a time. Error bars = 1 SD.

**Figure S1:**
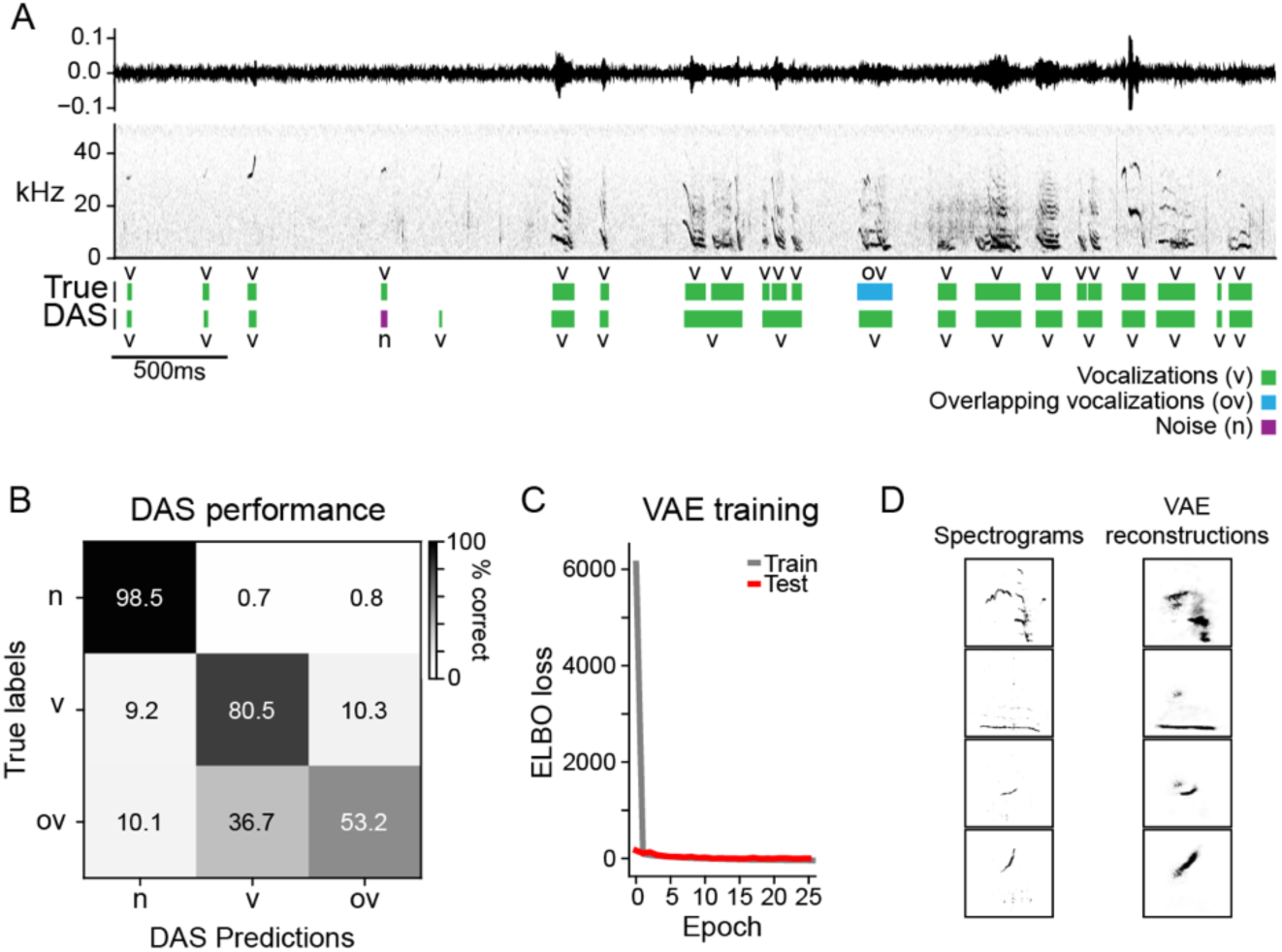
Unsupervised extraction of vocalization categories. **A.** Example 5 seconds of overnight audio recording. Top: waveform. Middle: spectrogram. Bottom: True (human-labeled) labels compared to trained DAS network’s labels for vocalizations (v; green), overlapping vocalizations (ov; blue), or noise (n; purple). **B.** Confusion matrix of true versus predicted DAS labels. **C.** Evidence lower bound (ELBO) loss for training the variational autoencoder (VAE) model on spectrogram images. **D.** Examples of VAE reconstructions (right) of spectrograms (left) from the 4 vocalization categories used in this study.

To visualize how vocalization acoustics were distributed across the four categories, we used a UMAP representation to project the latents that explained the most variance into a 2D space (**Fig 1C**). We quantified how accurately the vocalizations could be classified into these 4 categories by training a random forest classifier on the latent representations of the vocalizations (see STAR METHODS). Across 100 iterations of randomly sampled train and test sets, the classifier performed 96-98% on all four vocal categories (**Fig 1D**), indicating that family-invariant vocal categories can be described by their acoustics among hundreds of variants.

Although it is known that gerbils exhibit family-specific usage of vocalizations (Peterson et al., 2024), it has not been shown whether the acoustic properties of any category differs between families. To test whether vocalizations within each of the four categories can be classified by family-of-origin, we performed the classifier analyses described above, but trained the models for one vocalization category at a time and used family identity as the classifier labels. When trained on randomly sampled vocalization latents from all 5 families, family identity could be decoded with 35-43% accuracy, compared to 20% chance (**Fig 1E**, left). Across all neural experiments, two families were kept constant for all subject animals. When trained with just these 2 families, family identity could be decoded with >65% accuracy across all four vocalization categories (**Fig 1E**, right). These results indicate that gerbil family identity information is embedded within the acoustics of single vocalization categories, although this information is not as robust as category itself.

### Diverse AC neuron responses to gerbil vocalizations

Since the acoustic features of each vocalization can be classified based on category and family-of origin (**Figure 1**), we next asked how well they could be classified by AC neuron activity. A 64-channel silicon probe was implanted in the left AC of adult gerbils (*n*=5; see STAR METHODS) and, after recovery, wireless recordings were obtained from awake, freely moving animals while the stimulus set was presented from an overhead speaker (**Fig 2A**). As illustrated in **Figure 2C**, the stimulus set consisted of four vocalization categories, vocalization bouts, noise stimuli, and pure tones. For each vocalization category, variants from 3 families were presented, with one of the families always being the subject animal’s own family. Thus, each neuron was stimulated with ∼1500 unique variants. Vocalizations were presented in 5-minute blocks of the same category, with family identity of the variants randomly shuffled throughout these blocks. Each individual variant was presented 5 times.

**Figure 2:**
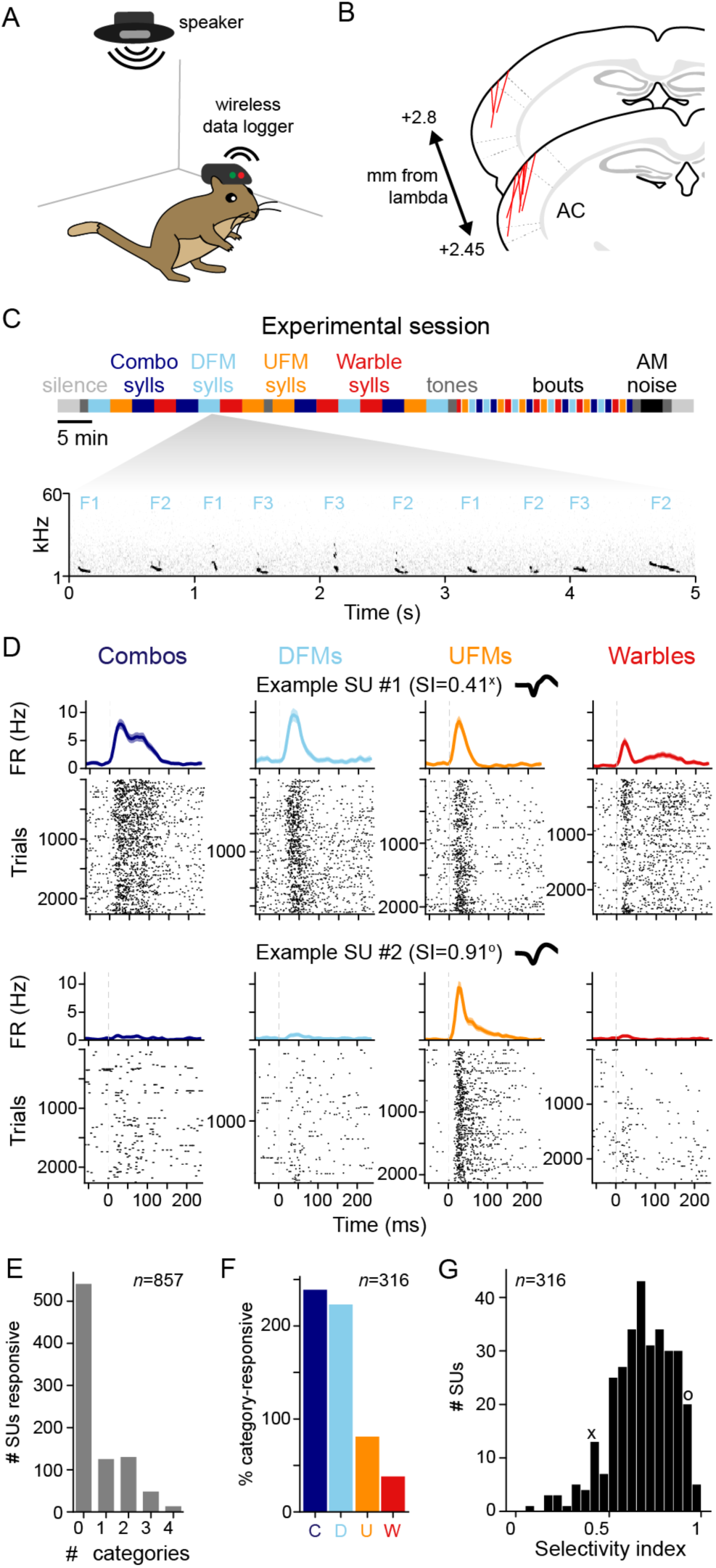
Heterogeneous responses of AC units to gerbil vocal categories. **A.** Schematic of gerbil implanted with wireless electrophysiology system. **B.** Overlays of probe tracts on the AC sections from the Radtke-Schuller stereotaxic axis. **C.** Stimulus presentation for the whole experimental session (F1, F2, & F3 represent Family ID 1, 2, & 3). **D.** Two example single units’ (SU) responses to all trials within a vocal category. Top: Mean peri-stimulus time histogram (PSTH) to all trials in a given vocal category (shading = 95% confidence interval). Bottom: Raster plot of spike times on every trial. Inset: Mean waveform of the single unit. **E.** Quantification of percentage of AC population that responds to increasing number of vocal categories. **F.** Quantification of percentage of responsive AC population that responds significantly to each vocal category. **G.** Distribution of selectivity index of all responsive units.

We recorded auditory responses from 857 AC single units across the 5 subject animals (*n*=53-299/animal; mean = 171.4, SD = 85.4), confirmed by referencing the electrode tracts to the gerbil stereotaxic atlas (**Fig 2B**; Radtke-Schuller 2016). We observed a range of single unit responses to the vocalization categories with some neurons responding reliably to all four categories (**Fig 2D**, top), and others showing high vocalization selectivity (**Fig 2D**, bottom). Within the entire population, 37% (*n*=316) of AC neurons exhibited a significant change in firing rate to at least one vocalization category (paired *t*-test between spontaneous and vocalization-evoked firing rates with Bonferroni correction; *p*<2.5e-3). “Category-selective” neurons that responded significantly to only one category comprised 15% (*n*=125) of the population (**Fig 2E**). The remaining 22% (*n*=191) of neurons responded to 2 or more categories. Of the units responsive to at least 1 category (“vocal-responsive units”), the majority of cells responded to the Combo and DFM vocalizations, with smaller portions of the population responding to UFMs and Warbles (**Fig 2F**). For these vocal-responsive units, a selectivity index (SI) was computed to quantify how specific or general category responses were. An SI of 1 indicates that the unit responded to only 1 category, while an SI of 0 indicates the unit responded equally to all 4 categories. The two example units in **Fig 2D** displayed SI values of 0.41 and 0.92. The median SI for the entire vocal-responsive population was 0.72 (**Fig 2G**), suggesting that vocalization-responsive AC neurons were largely category selective.

We next asked whether AC neurons responded to family-specific acoustic features and could plausibly contribute to vocal identification. Many single AC neurons were modulated by family identity for one or all vocal categories tested (**Fig 3A-B**). Of the category-responsive units, 22% (*n*=68) of neurons’ firing rates were significantly modulated by family identity for at least one vocal category (one-way ANOVA with Bonferroni correction; *p*<2.5e-3). For each implanted animal, we presented vocalizations from its own family and from 2 other families. We found that the proportion of AC neurons that was responsive to own-compared to other-family was not significantly different for any vocalization category (**Fig 3C**), suggesting that family experience does not upregulate the number of own-family-responding units. Further, the median selectivity index for family identity was low, ranging from 0.24-0.29 across vocal categories (**Fig 3D**).

**Figure 3:**
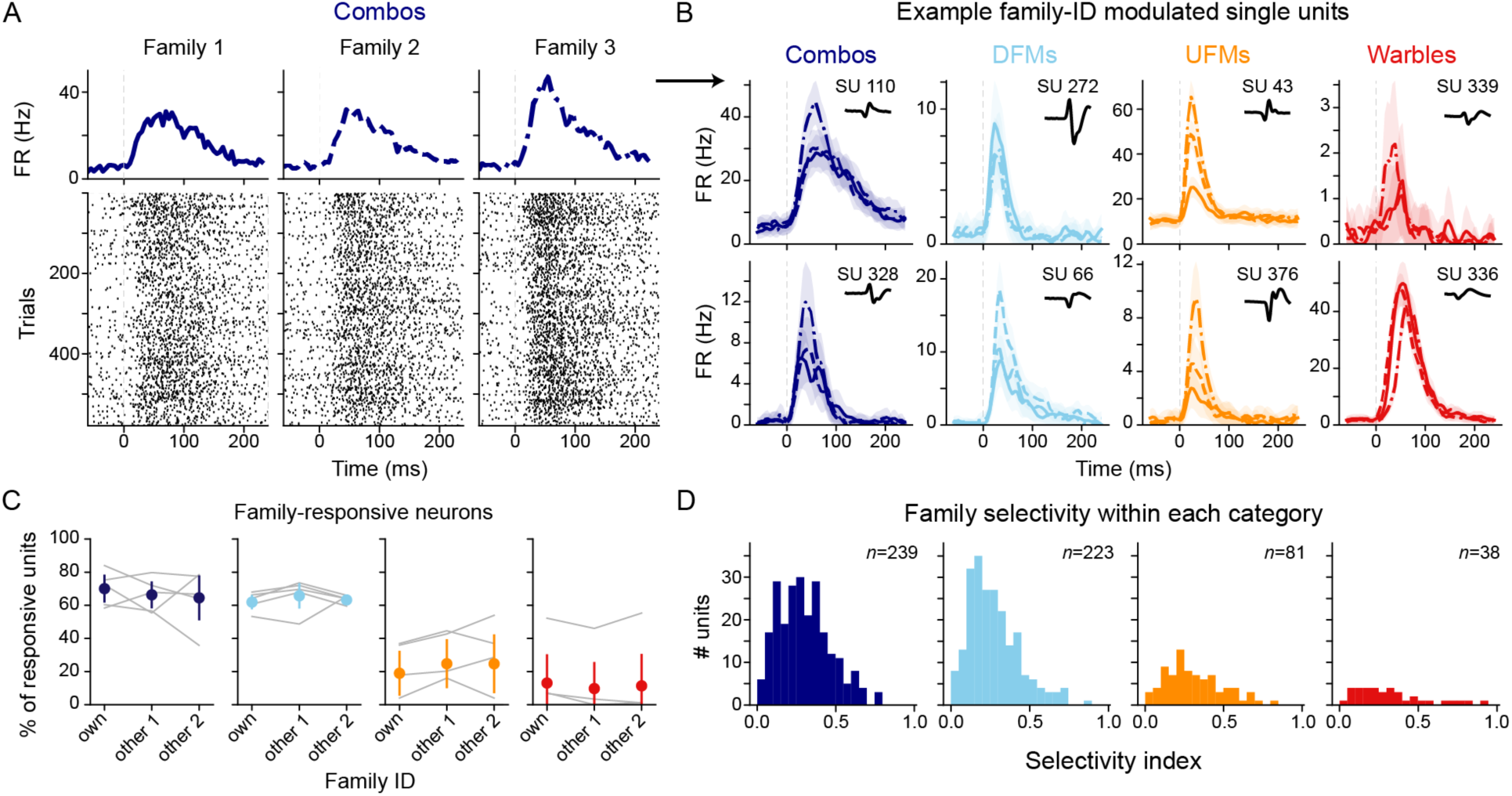
Single units in AC are modulated by family identity of vocalizations. **A.** Example single unit response to all trials of Combos produced by 3 gerbil families. Top: Mean peri-stimulus time histogram (PSTH) to all trials in a given vocal category. Bottom: Raster plot of spike times on every trial. **B.** Eight example single unit average PSTHs to all variants in a vocalization category, separated by which family produced the variants (shading = 95% confidence interval). **C.** Percent of population that is responsive to own v other family for each vocalization category. Colored points and error bars = mean and 95% confidence interval across all animals. Grey lines = individual animals. **E.** Selectivity index of category-responsive cells to family identity.

We also characterized AC neuron responses to arbitrary waveforms (tones, noise) and asked whether they could explain the observed vocalization-evoked responses. In addition to the vocalization variants, each neuron was stimulated with pure tones (30 ms, 1.5-50kHz) and amplitude-modulated (AM) noise (depth: 100% depth, rates: 2-64Hz; passbands: 1-10, 1-25, 5-25, 25-40, or 30-55 kHz). 43% of AC units (*n*=367) were responsive to either vocalizations, tones, or AM noise (**Fig S2A**). Of these sound-responsive neurons, 86% were responsive to vocalization-responses, indicating that AC neurons may be more easily driven by vocalizations than artificial stimuli. We found neurons that significantly responded to frequencies spanning the entire frequency range of the vocalization stimulus set (**Fig S2B-C**), indicating that we adequately sampled the tonotopic axis. To compare vocal and non-vocal responses, we computed the phi correlation (a value of 1 is a perfect positive correlation) between the presence of a vocalization response for each category and the presence of a tone or AM response for matched frequencies (**Table 1**). Although we found small to moderate correlations for all categories (0.12-0.64), this result suggests that vocalization responses are not explained by frequency or modulation tuning alone.

**Supplementary Figure 2:**
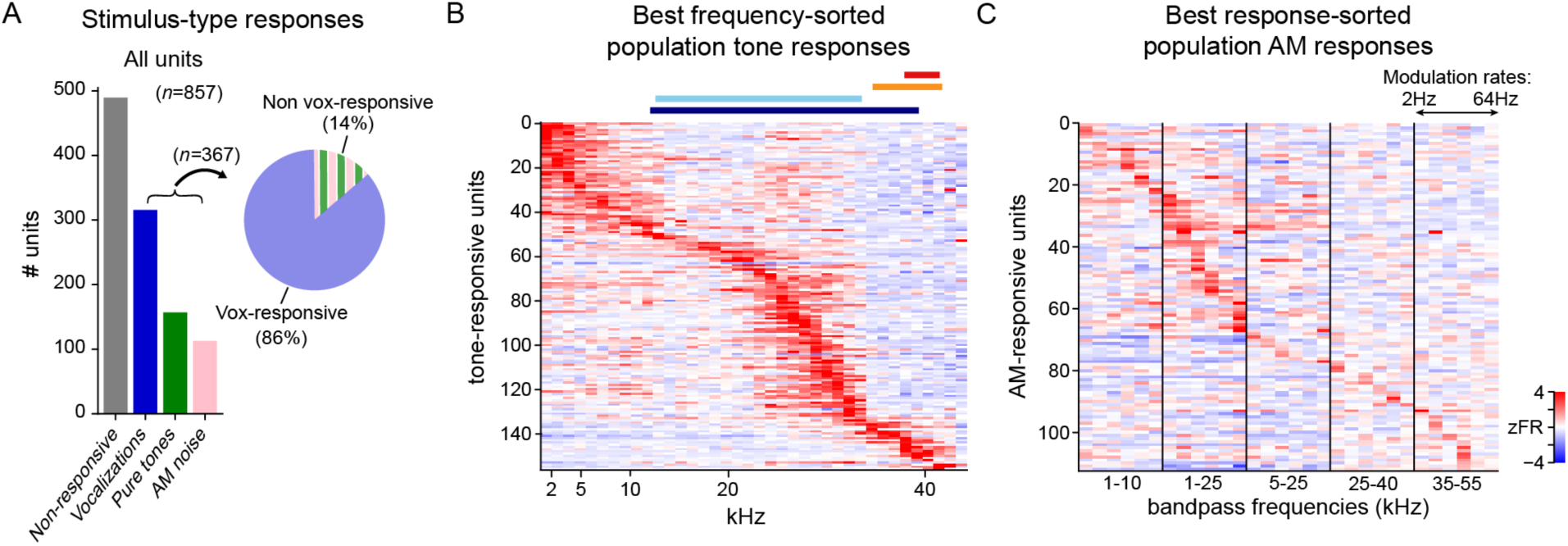
Single units responses to vocalizations, tones, and noise. **A.** Distribution of single units that were significantly responsive to vocalizations, tones, AM noise, or none. **B.** zFR response to tones for all tone-responsive units, sorted by their best frequencies. Colored bars above the heat map correspond to the frequency content of the 4 vocalization categories. **C.** AM-noise responses sorted by their best bandpass frequency and modulation rate.

**Table 1:**
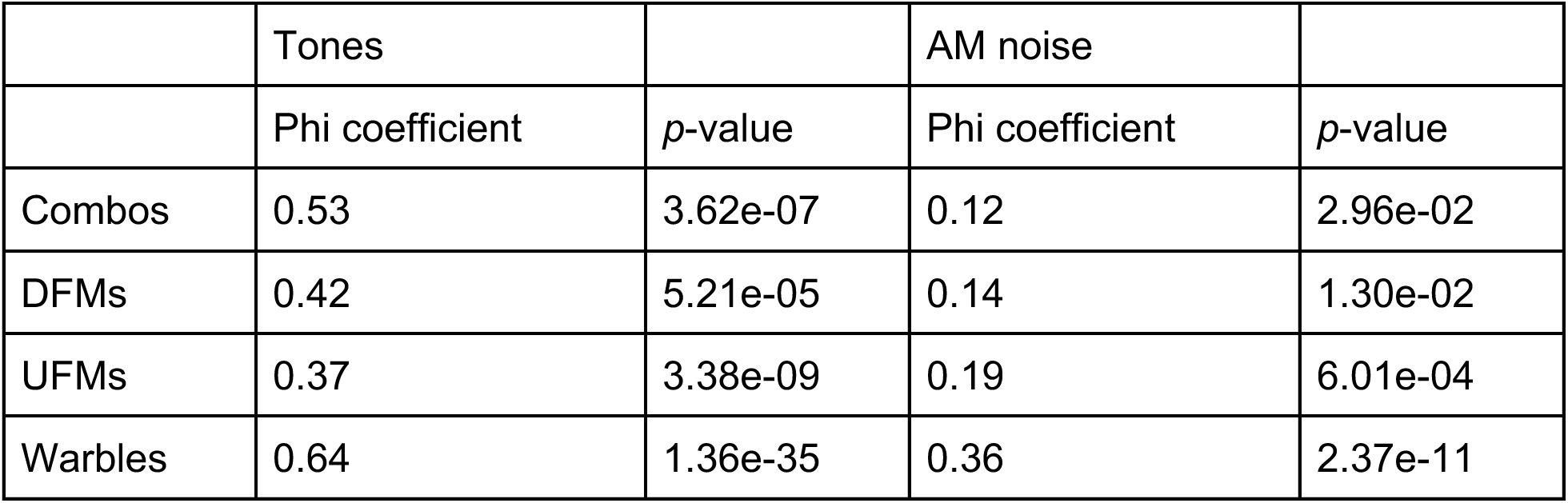

|  | Tones |  | AM noise |  |
| --- | --- | --- | --- | --- |
|  | Phi coefficient | p-value | Phi coefficient | p-value |
| Combos | 0.53 | 3.62e-07 | 0.12 | 2.96e-02 |
| DFMs | 0.42 | 5.21e-05 | 0.14 | 1.30e-02 |
| UFMs | 0.37 | 3.38e-09 | 0.19 | 6.01e-04 |
| Warbles | 0.64 | 1.36e-35 | 0.36 | 2.37e-11 |

Since animals were freely moving, their position could change relative to the speaker during the course of testing. To determine whether positional factors were a source of AC neuron response variance, we asked how every tone-responsive neuron’s z-scored firing rate was modulated as a function of the animal’s distance to the speaker, head orientation to the speaker, velocity, or acceleration (**Fig S3**). The 5th and 95th percentiles for the slope fit R^2^ values were 0.004 to 0.25 across the four behavioral variables, with only 2 neurons having slope fits that reached significance (Pearson correlation with Bonferroni correction; *p*<2.5e-3), suggesting that movement and position had a relatively small influence on neuronal responses.

**Supplementary Figure 3:**
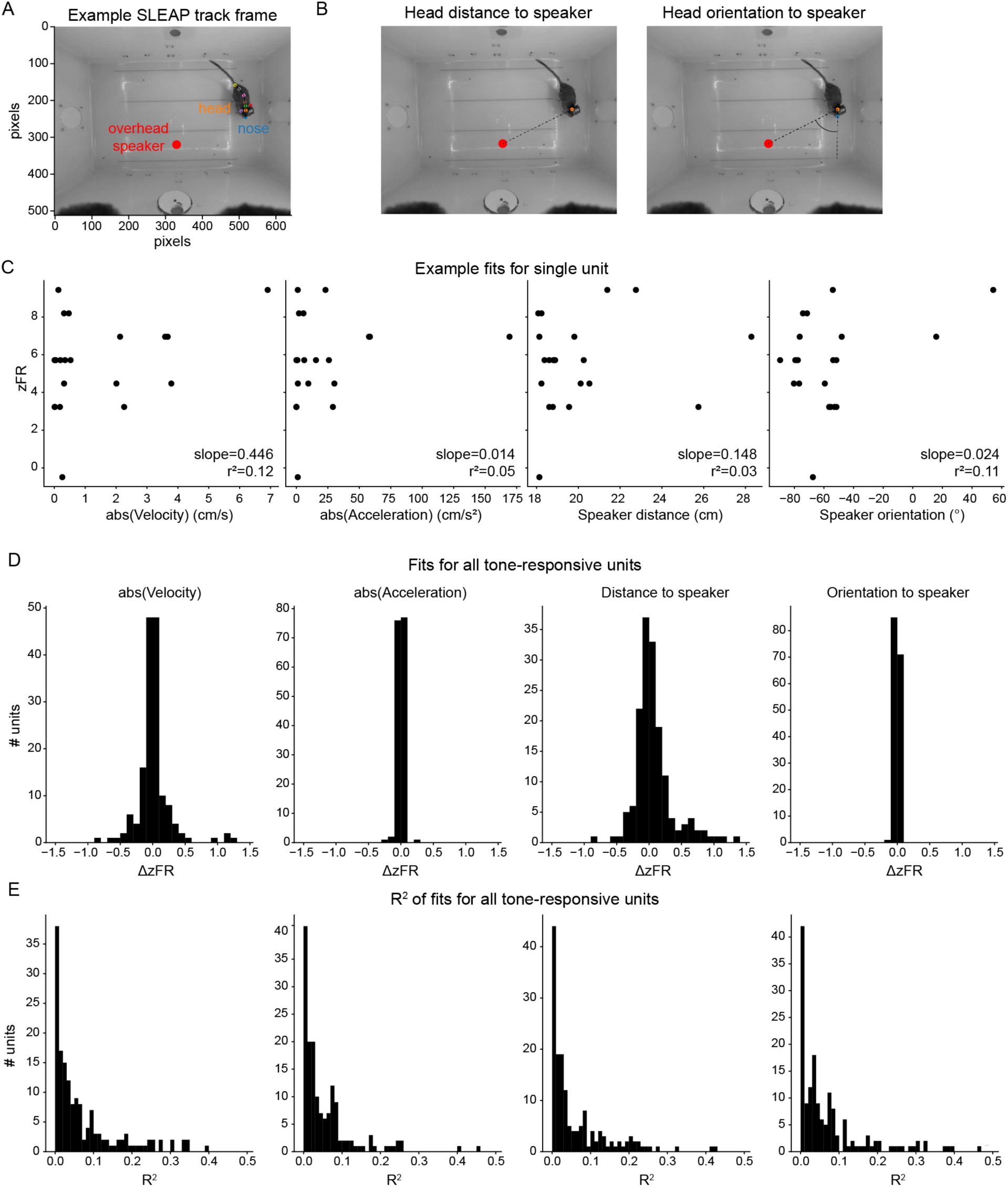
Single units modulation by movement variables. **A.** Example frame showing SLEAP tracks of animal and position where the overhead speaker points to. **B.** Diagrams demonstrating calculation of distance to point directly below the speaker and head orientation relative to the speaker. **C.** Example scatter plots of the 4 movement variables versus zFR for the single unit’s best frequency. **D.** Distributions of fit slopes for all tone-responsive units in the population to the 4 movement variables. **D.** Distributions of fit R^2^ for all tone-responsive units in the population to the 4 movement variables.

### Vocalization category and family identity can be simultaneously decoded from AC populations

Since single units were modulated both by vocal category and family identity, we next asked how well they could be decoded from AC population activity. Since many units demonstrated heterogeneous temporal response patterns, we built a linear support vector machine (SVM) decoder using single trial, z-scored 30ms PSTHs. **Figure 4A** provides an overview of the z-scored PSTHs for the vocal-responsive pseudopopulation, with all PSTHs for a single category averaged for each neuron. Therefore, this visualization does <u>not</u> capture the variant-to-variant differences in PSTHs that are present within our test data (**Fig 2D, 3A, S4A-B**). To compare vocalization category and family identity decoding, these analyses were performed in parallel such that the same trials were used for each decoder. Single-trial PSTHs (250 trials per family per category) for the entire population were randomly sampled and split into train and test sets. The vocalization category SVM is trained to form 4 linear hyperplanes that best separate the population activity for one category from the activity of the 3 other categories (**Fig 4B**, top). In order to perform family identity decoding for our pseudopopulation, we had to restrict the analysis to the 2 families that all animals had in their stimulus set (since ‘own’ family was not the same for all animals). Therefore, the family identity SVM forms one hyperplane that best separates Family 1 and Family 2 within the same vocal category (**Fig 4B**, bottom). This procedure is performed 100 times with new subsamples of the trials. Note that the decoders performed either equally or better when restricting the population to only the vocal-responsive units (**Fig S4C-D**), so only vocal-responsive units were used in subsequent analyses. Further, the PSTH decoders always outperformed a simple firing rate decoder (**Fig S4E-F**).

**Figure 4:**
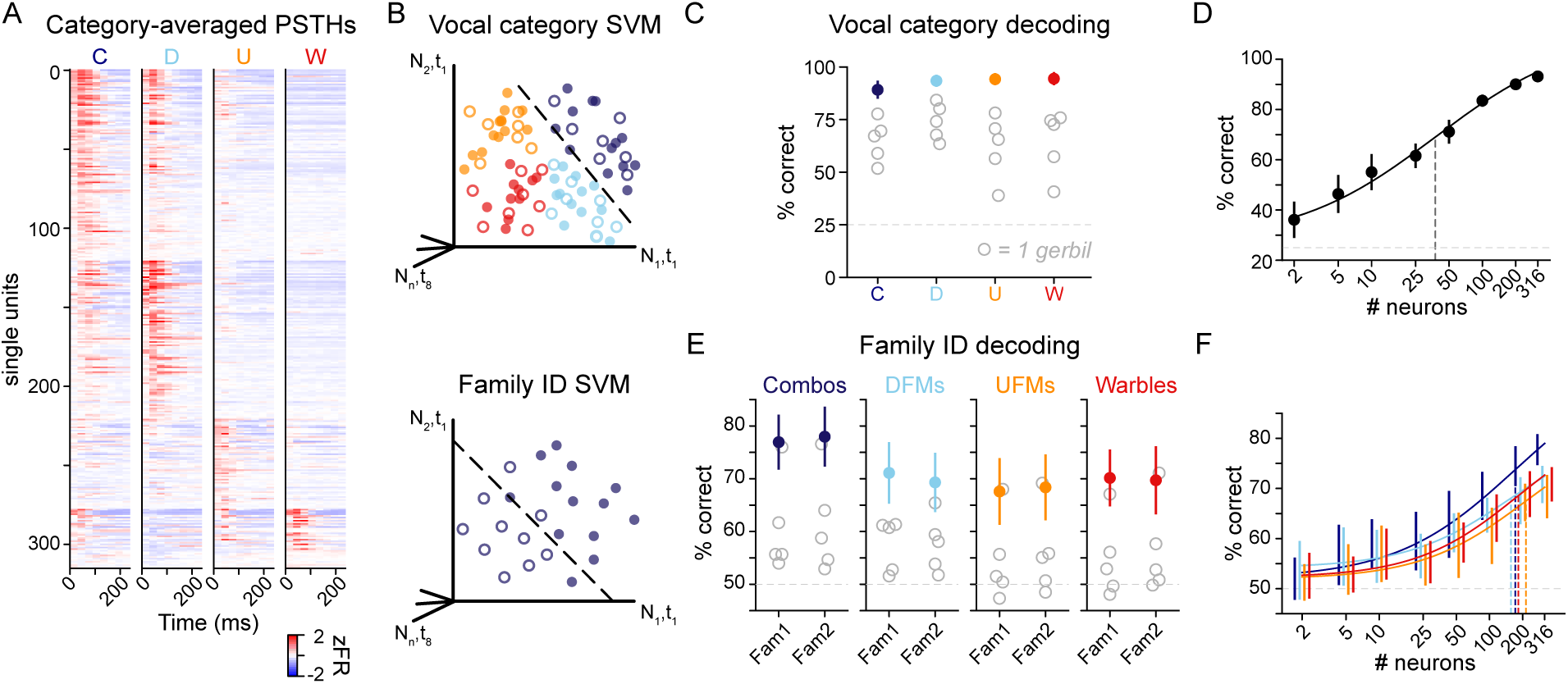
Vocalization categories and family identity can be decoded from AC population activity. **A.** Category-averaged 30ms-bin PSTHs (zFR = z-scored firing rate) for every vocal-responsive neuron (*n*=316). Neurons are sorted by the vocalization category they showed the largest response to. **B.** Schematic of linear SVM classifiers (top: vocalization category, bottom: family ID for 1 category at a time). **C.** Vocalization category SVM performance (*n*=100 iterations resampling 250 trials from each category). Colors = pseudopopulation performance (*n*=316 neurons). Grey circles = individual gerbil performance. Error bars = 1 standard deviation (SD) across 100 iterations. Chance = 25%. **D.** Vocalization category SVM performance versus number of neurons sampled from the population. Error bars = 1 SD across 20 iterations. Solid line = sigmoidal fit to the means. Dashed line = inflection point of the fit. **E.** Family ID SVM performance (*n*=100 iterations resampling 250 trials from each family, within a category). Colors = pseudopopulation performance (*n*=857 neurons). Grey circles = individual gerbil performance. Error bars = 1 SD across 100 iterations. Chance = 50%. **F.** Family ID SVM performance versus number of neurons sampled from the population, for each vocal category. Error bars = 1 SD across 100 iterations. Solid line = sigmoidal fit to the means. Dashed lines = inflection points of the fit.

**Supplementary Figure 4:**
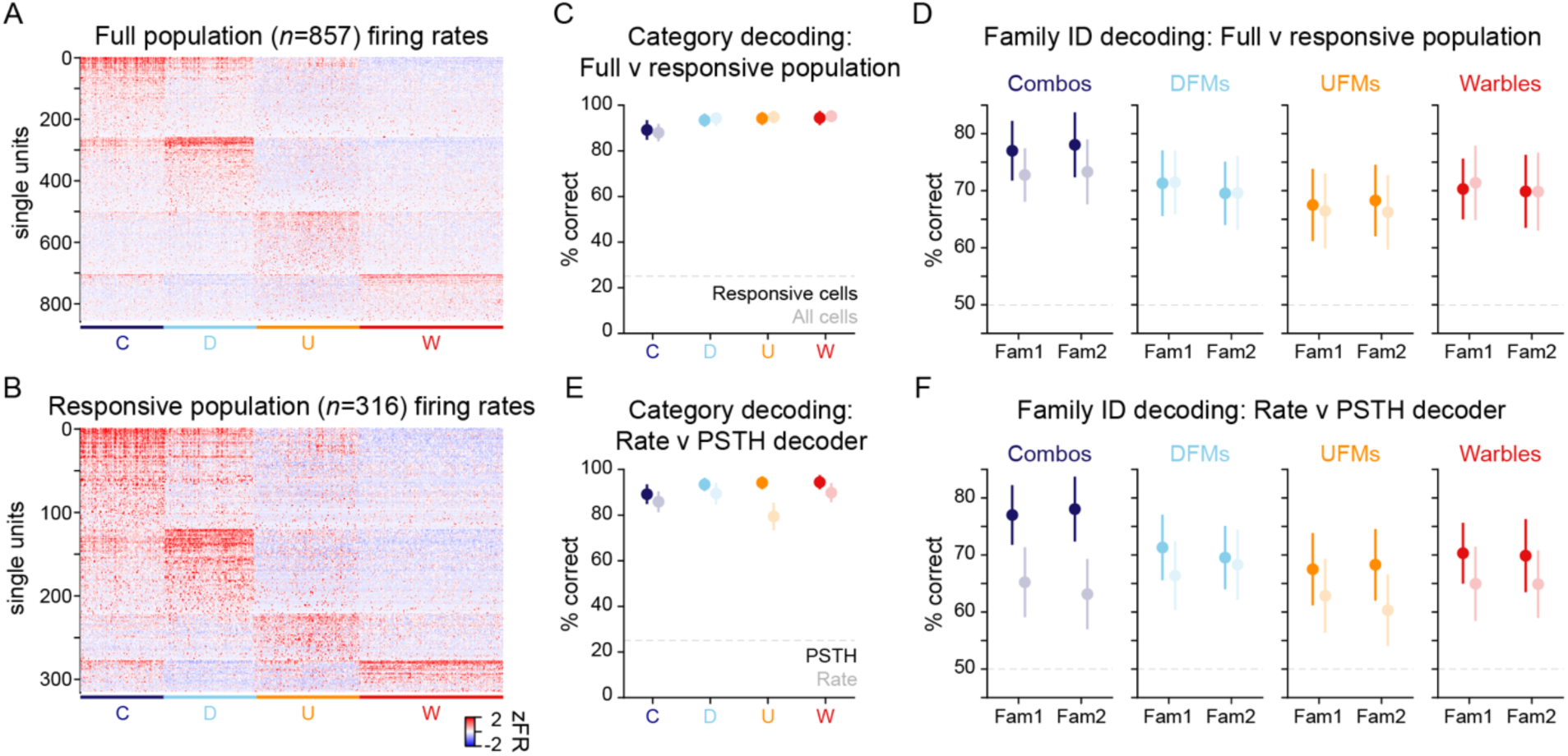
Responsive-unit, PSTH decoders outperform full-population and rate decoders. **A.** z-scored firing rate matrix for the entire population for all trials in the experiment, sorted by category (1 column = 1 trial). **B.** As in A, but restricted to the *n*=316 responsive units. **C.** Performance of vocalization category SVM using either the entire population (lighter colors) or just the responsive units (darker colors). Error bars = 1 SD across 100 iterations. **D.** As in C, but for family ID decoders. **E.** Performance of vocalization category decoder using the z-scored firing rates (lighter colors) or the z-scored 30ms PSTHs (darker colors). **F.** As in E, but for family ID decoders.

**Supplementary Figure 5:**
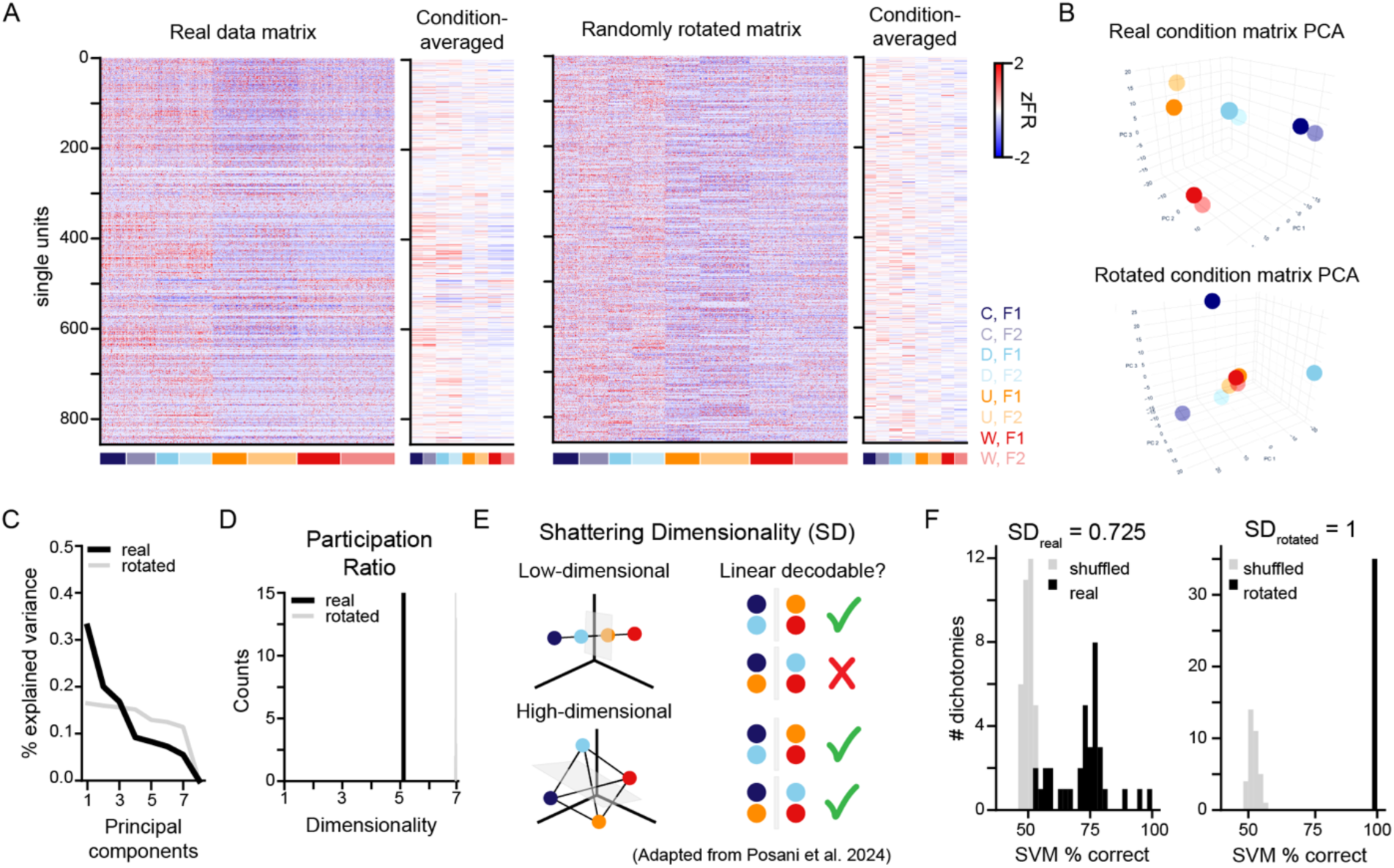
AC populations represent the natural stimulus set in a lower dimensionality than is expected by chance. **A.** Left: Z-score firing rate matrix for all units in the population. A single column represents a neuron’s mean z-scored firing rate to all trials of a given variant. Columns are grouped by vocalization category and family ID condition (colored along the bottom; labels to the right). Right: Same matrix as on the left, but the row order has been shuffled independently for each vocalization category/family condition. Condition averaged matrices show the same data, but all firing rates for variants within a condition are averaged. **B.** 3D PCA plots of the true versus rotated matrices. **C.** Explained variance of the principal components for the true and rotated matrices. **D.** Participation ratio of the real data (black line) versus 100 random rotations of the data (grey). **E.** Schematic demonstrating intuition for shattering dimensionality. **F.** Shattering dimensionality of the real versus rotated data for SVMs trained with the true trial labels (black) versus shuffled (grey).

We found that both vocalization category and family identity could be decoded from AC activity significantly above chance, despite being tested with hundreds of unique vocalization variants. Vocalization category performance was 89-94% across the 4 categories (**Fig 4C**; one-sample *t*-test *p*<4.05e-163), close to the upper limit defined by the acoustic classifier in Figure 1D. Even in individual gerbils, with responsive neural sample sizes of 25-89 single units, performance was significantly above chance, demonstrating robust vocal category representations by AC populations. There were differences in performance across the four vocalization categories, with performance ranging from 89% for the Combos to 94% for the Warbles. Combos may be more difficult to classify because they include acoustic features from DFMs and Warbles. We further asked how many neurons are required to perform vocalization category classification by subsampling the pseudopopulation with increasing population sizes (**Fig 4D**). We fit the mean performance with a sigmoidal function to find how many neurons were required for 50% of maximal performance (the inflection point). We found that 37 neurons are required to perform 62% correct, a relatively small population size given the variance of the stimulus set.

Across all 4 vocalization categories, the AC population classified family identity with 68-78% accuracy, significantly above 50% chance (one-sample *t*-test *p*<1.5e-65; **Fig 4E**). This is comparable to decoding of family identity from the latents that represent individual vocalization acoustic features (**Fig 1E**, right). For most vocalization categories, family identity could be decoded from an individual gerbils’ AC activity. Family identifying features were present in all of the vocalization categories, although the SVM performed best for Combo vocalizations. The population size needed to perform family ID decoding across vocalization categories ranged from 167-178 units (**Fig 4F**), around 5 times as many needed for decoding vocalization category, suggesting that this is a more difficult feature for AC neurons to extract from individual vocalizations.

Given that AC populations can both generalize within vocalization category and discriminate between different families, we further assessed the dimensionality of the population representations which can have implications for cognitive flexibility and memory capacity (Posani et al., 2024; Rigotti et al., 2013). For example, a low-dimensional representation, where the only variance in the population is along the coding-axis for category, would allow AC to perfectly discriminate between vocalization categories. However, if there is no other variance in the population, then the AC would be unable to discriminate between different families within a class. A maximally high dimensional representation would allow AC to discriminate between any pair of vocalization categories and family ID combination. Dimensionality was assessed with the participation ratio and shattering dimensionality of our dataset. The participation ratio quantifies the dimensionality based on how distributed variance is across all dimensions of the data (e.g., a value of 1 would be minimally dimensional and 7 would be maximally dimensional for a mean-centered, 8-condition dataset). We found that our data had a participation ratio of 5.1 (**Fig S5C-D**), indicating that AC populations indeed compress vocalization representations into a lower dimensional space than is expected from a random geometry. The shattering dimensionality measures the mean performance of linear decoders on all possible dichotomies of the data (**Fig S5E-F**). A value of 0.5 indicates the lowest dimensionality, as the decoders did not perform above chance across all dichotomies, while a value of 1 would indicate maximal dimensionality, as all dichotomies are linearly separable. We calculated a shattering dimensionality of 0.725, again demonstrating that the AC population has a low dimensional geometry that may underlie generalization across variants to represent vocal categories.

### Individual AC neurons contribute to category and identity decoding

An advantage of the stimulus paradigm is the inclusion of both category and identity variance, permitting us to ask whether single AC neurons are specialized for the extraction of one type of information. To address this question, we reran the SVM 1000 times to compare the contribution, or relative weight (**Fig 5A**), of each neuron to the population decoders. We compared the contribution of each neuron to the vocalization category and the family identity SVMs to their weights in a null decoder trained with shuffled trial labels. A null decoder will equally weigh all neurons in the population across 1000 iterations, since no particular neurons give the null SVM more information to perform classification. Thus, this analysis compares the contribution of neurons relative to a perfectly distributed population code (all neurons contributing equally). A paired *t*-test (*p*<2.5e-3) was used to compare the true and null contribution for all family decoders and for each category, the results of which we used to label each cell as strongly contributing only to category, only to family identity, to both, or to neither decoder (**Fig 5B**).

**Figure 5:**
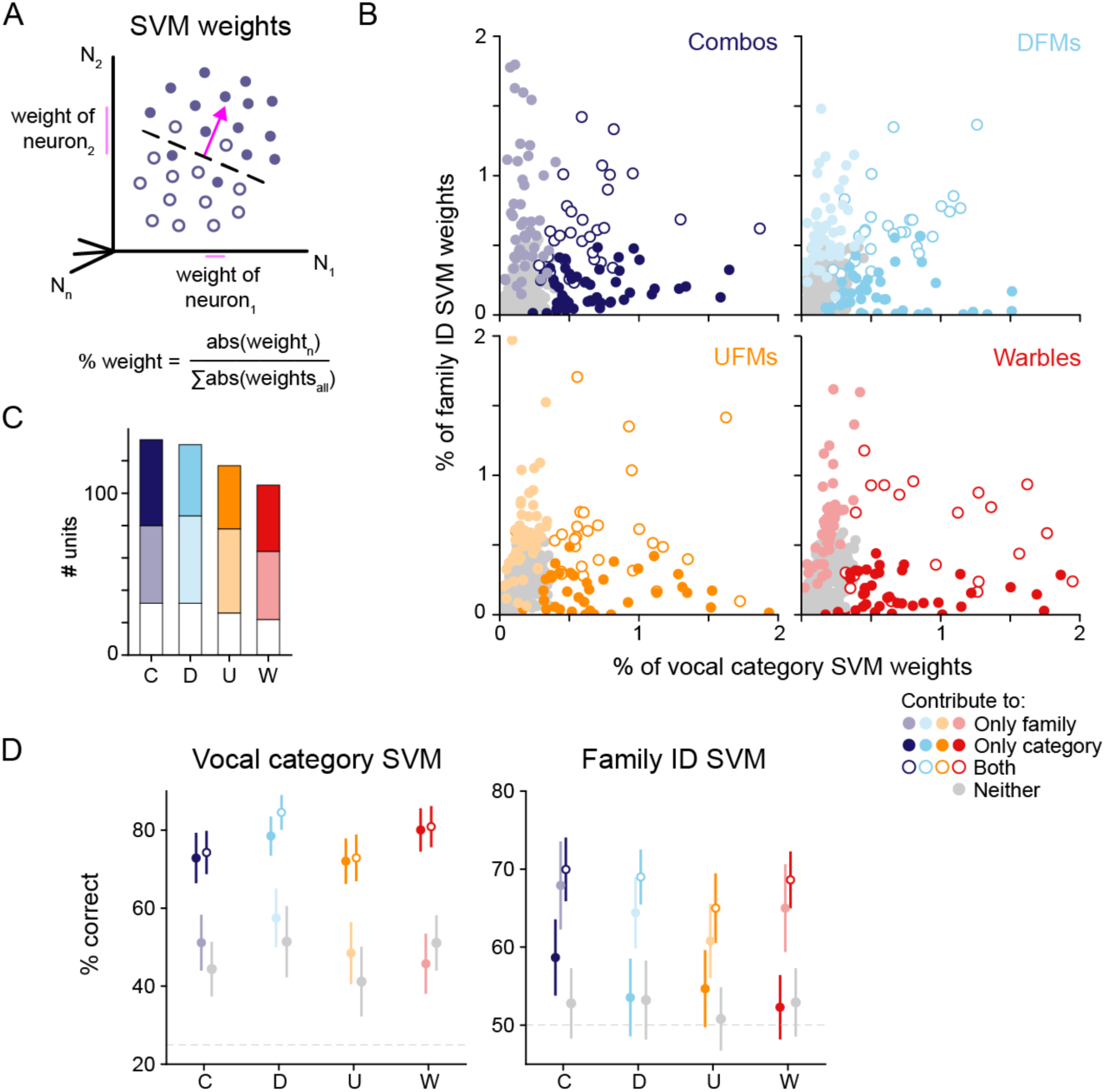
AC neurons contribute to both category and identity decoding. **A.** Schematic demonstrating what is meant by an SVM weight. **B.** Absolute values of SVM weights for all responsive units (*n*=316), shaded by whether they contribute to that vocalization’s category classification, family identification, both, or neither. **C.** Quantification of the number of units contributing to either or both, or neither of the classifiers described in B. **D.** Performance of SVMs only using 20 sampled neurons from the category only, family only, both, or neither populations.

There were more responsive cells that contributed significantly to *either* the vocalization category or the family identity decoder than to both decoders (**Fig 5C**), suggesting the participation of ‘specialists’. However, the ‘generalists’ (neurons that contributed significantly to both decoders) performed classification just as well as the specialists (**Fig 5D**). This suggests that single neurons can read out information to more than one downstream decoder without impairing either decoder’s performance. We found that vocalization category can be decoded from ‘family only’ and ‘neither’ cells at ∼50% when subsampling 20 neurons of each neuron type (**Fig 5D**), suggesting that even though these neurons don’t contribute strongly on their own, vocalization category can be extracted from a purely distributed code at the population level. Since family identity could not be decoded from ‘only category’ or ‘neither’ cells, this suggests that family identity cannot be extracted from a purely distributed code, but rather a sparse code with a few strongly contributing neurons. The exception was Combo vocalizations, where family ID could be decoded significantly above chance from the ‘category only’ cells (1-sample *t*-test, *p*=2.75e-7), suggesting again that the combinatorial nature of these vocalizations offers the population more information from which to extract both categorical and social information.

### AC representation of vocalization categories displays a boundary effect

We have thus far shown that even in the presence of many variants and identity variation, vocalization category can be decoded from AC population activity. To determine whether the AC is forming discrete group representations (as opposed to simply discriminating), we asked whether neurons displayed a boundary effect, wherein discrimination across the boundary separating two categories is greater than discrimination within a category ((Holt & Lotto, 2010); **Fig 6A**). We addressed this issue by directly comparing the ability of an SVM to decode vocalization category either from the z-scored PSTHs or from the vocalizations’ acoustic features. For acoustic features, we used either the latents or specific extracted measures such as frequency, bandwidth, duration, etc. (see STAR METHODS). We randomly subsampled groups of 25 variants either from the same vocalization category (within-category) or from different vocalization categories (across-boundary) and asked an SVM to perform binary classification between the 2 groups (**Fig 6B**). If the AC representations exhibit a boundary effect, we would expect either greater neural across-boundary classification or poorer within-category classification, as compared to the acoustics.

**Figure 6:**
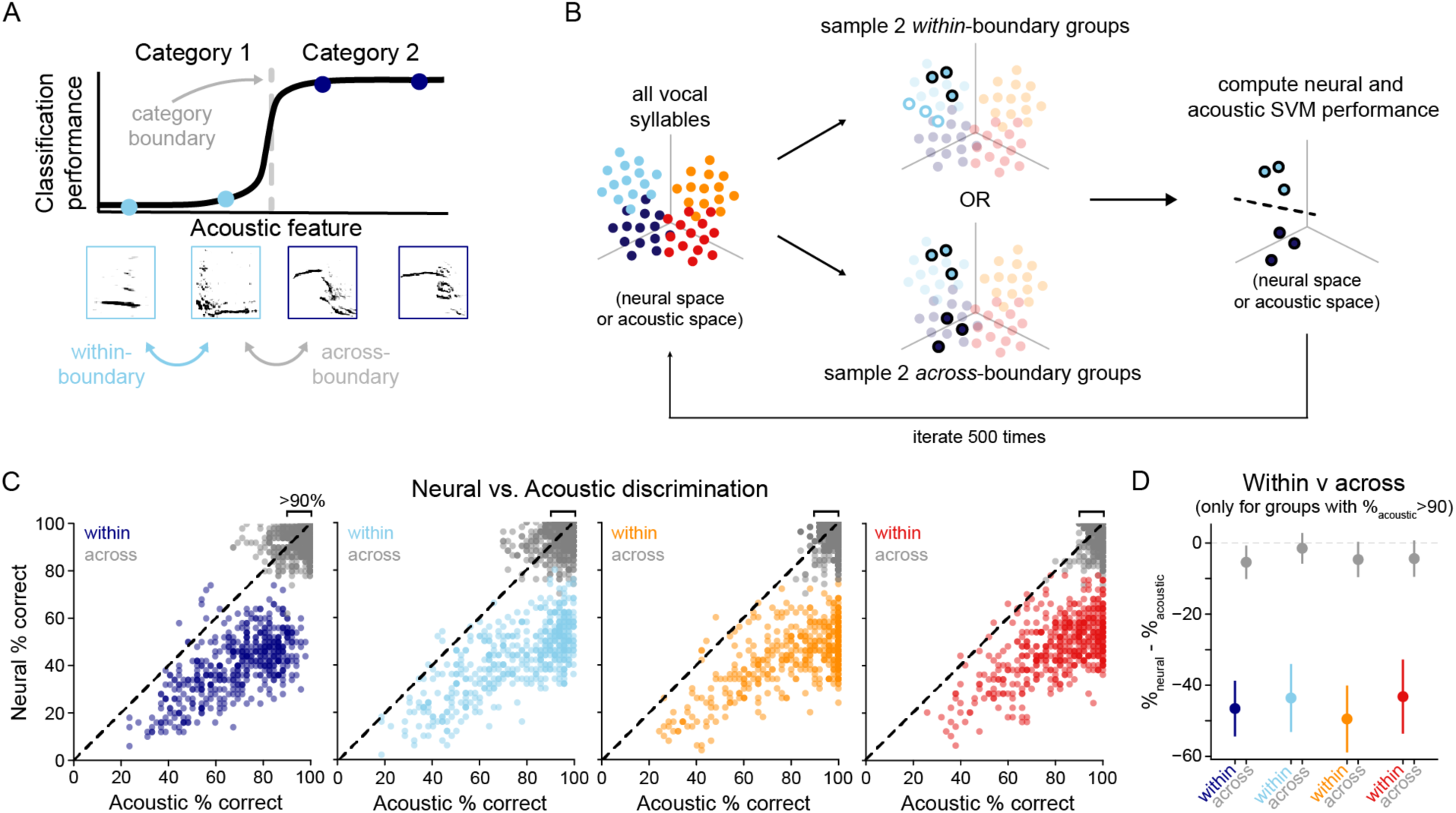
AC represents the vocalization as categories. **A.** Top: schematic of the categorization boundary effect (Kronrod et al., 2016). Bottom: example spectrograms illustrating a within-boundary vs. across boundary pairwise comparison. **B.** Method schematic for directly comparing acoustic and neural decoding of groups of variants within-vs. across-category boundaries. **C.** Acoustic versus neural SVM performance across 500 iterations of sampling 2 groups of 25 variants. Colors = within-category iterations, grey = across-category iterations. **D.** Neural - acoustic performance for iterations where acoustic performance was >90%.

We found that classification was poorer within-category as compared to across-boundary, both for the neural and the acoustic SVMs (**Fig 6C**: gray symbols, across boundary; color symbols, within-category). We then restricted our analysis to only those variants for which the acoustic SVM performed within-category classification at >90% (**Fig 6D**). Here, we found that the neural SVM performance was effectively at chance (∼45%). Therefore, the AC represents vocalization categories not as direct manifestations of their acoustic features, but rather as discrete groups.

## DISCUSSION

Vocalization processing involves two distinct requirements: generalizing across many voices to categorize a sound’s meaning and, inversely, distinguishing between many voices to recognize identifying information. In humans, both meaning and identity information can be extracted from vocalizations within 300 ms of sound onset (Allopenna et al., 1998; Grosjean, 1980; Lavan & McGettigan, 2023; Mila Mileva & Nadine Lavan, 2023; Owren et al., 2007), consistent with simultaneous extraction of meaning and speaker identity (Chandrasekaran et al., 2011). At the neural level, these two processes have been explored in separate studies that typically avoid the impact of vocalization variability. Here, we study generalization and identification concurrently using a >1500 stimulus set of gerbil vocalizations belonging to four vocal categories and five gerbil families while recording single unit responses in the AC of awake adults.

While a neural representation of vocalization categories has been characterized in several species (Montes-Lourido et al., 2021b; Prather et al., 2009; Robotka et al., 2023), these studies used relatively few variants and could not determine whether the representation was robust to the natural range of variance. However, several studies have demonstrated neural categorization by presenting synthetic variants that were morphed from a small set of natural vocalizations (Carruthers et al., 2015; Holmstrom et al., 2010; Lawlor et al., 2025; Prather et al., 2009; Town et al., 2018; Tsunada et al., 2011). These studies show nonlinear neural transformations of vocalizations into categories across an artificially manipulated acoustic axis, and imply a similar transformation for a natural range of variants. However, morphing could plausibly introduce unnatural feature combinations that would not be generalized in natural vocalizations. Finally, a few studies have shown that AC neurons can categorize a few dozen natural variants of phonemes or vocalizations (Mesgarani et al., 2008; Shepard et al., 2015), although these studies did not challenge the neural populations on both category and identity classification. Taken together, these studies provide deep insight into the neural basis of vocalization categorization, while leaving open the question of how social identity cues are represented simultaneously within categories.

Our study confirms and extends these results by challenging each AC neuron with ∼1500 natural variants that derive from 4 vocalization categories and 3 separate gerbil families (**Fig 1**). Using this approach, we demonstrate that decoding of both vocalization category and family identity is robust to natural acoustic variability. Consistent with previous reports, we found a sparse representation of vocalization-responsive neurons. However, the 316 vocalization-responsive AC neurons were sufficient to classify single trials into the 4 vocalization categories at a performance of 89-94% (**Figure 4C**), which is comparable to classification performance based on acoustic features (**Figure 1D**). Although this performance is higher than reported for classification of dozens of human phonemes by ferret AC (Mesgarani et al., 2008), this is likely explained by ferret study using more vocalization categories. Similar to a report on zebra finch auditory pallium (Robotka et al., 2023), we find that a small number of cells is required to perform categorization, perhaps due to the gerbil AC population having relatively strong category-selectivity at the single neuron level (median SI = 0.72; **Figure 2H**).

Perceptual categories can be defined operationally as the non-linear transformation of a continuous feature axis into distinct groups, separated by a sharp boundary across which discrimination of variants is greater than discrimination of within-category variants (Holt & Lotto, 2010; McMurray, 2022). This concept was originally demonstrated for human speech sounds (Liberman et al., 1957, 1961), and it has since been extended to nonhumans (Comins & Gentner, 2013; Elie et al., 2025; Green et al., 2020; Kar et al., 2022; Nelson & Marler, 1989; Seyfarth et al., 1980). For example, Elie et al. demonstrate that zebra finches can discriminate vocal categories across many individual vocalizers in a manner that is better explained by the vocalization’s behavioral usage than its acoustics. Here, we show that the AC population forms efficient representations of vocalization category in the presence of considerable identity variance (**Figure S5**), and performs within-category classification more poorly than would be expected from the acoustics (**Figure 6**). This result is reminiscent of a perceptual magnet effect (Kuhl, 1991) that may subserve category perception.

Behavioral evidence from many species suggest that certain vocalizations signal individual or colony-specific identity (Barker et al., 2021; Elie & Theunissen, 2018; Gerald G. Carter et al., 2008; Perrodin et al., 2020; Szenczi et al., 2016), but the neural mechanisms supporting this behavior remain uncertain. AC neuron responses can be tuned to conspecific vocalization features in squirrel monkey, marmoset, cat, and macaque (Perrodin et al., 2011; Wang & Kadia, 2001; Wollberg & Newman, 1972) and these responses have even been shown to be modulated by familiarity with a vocalizer (Agarwalla et al., 2023; Menardy et al., 2012). However, identity-specific information may differ across vocal categories (Elie & Theunissen, 2018), and representing identity would require AC neurons to parse subtle, high-dimensional, acoustic features on a category-by-category basis.

Here, we report that family identity can be decoded from an AC population for each of the vocalization categories (**Figure 4E**), suggesting that social identity is represented independently and concurrently within each category. This finding is consistent with a recent behavioral study showing that zebra finches can perform social discrimination from vocalizations across their repertoire, and that the features used for identification differ for each vocalization category (Elie & Theunissen, 2018). We show that family identification requires a larger AC population than does vocalization categorization (**Figure 4F**), suggesting this information is not as strongly encoded at the level of single variants and thus requires more neurons to extract this information. One limitation of our design is that single syllables were used as stimuli, whereas gerbils (and other rodents) typically emit vocalizations in a bout of several seconds (Peterson et al., 2024). Therefore, it is possible that vocalization repetition provides stronger social identity information, and temporal integration by a downstream decoder would be able to extract family-specific sound cues and perform better classification. Indeed, female zebrafinch neurons in NCM, a higher auditory area, respond preferentially to their mate’s full song compared to familiar and unfamiliar songs (Menardy et al., 2012) and bout responses, but not syllable responses, are enhanced in female AC after 3 days of male social experience (Agarwalla et al., 2023). A separate possibility is that primary AC serves to extract family-specific features and projects to a downstream decoder that compares those features to a family prototype, subserving ‘recognition’. This would be consistent with cognitive models positing that human voice recognition is a distinct mechanism from discrimination, given dissociations of these deficits in patients using lesion case studies (D. Van Lancker & Kreiman, 1987; D. R. Van Lancker et al., 1988; D. R. Van Lancker & Canter, 1982). While our findings support an AC mechanism for family identification, this interpretation is limited by the absence of behavioral measures for family recognition or categorization. Therefore, it is necessary to develop a paradigm that measures gerbil discrimination of vocalization categories and family-of-origin identities such that results can be compared to the neural responses reported here.

The neural computations underlying categorization of naturally-variable vocalizations are not well understood. If all the variance is distributed along the category axis, then the AC would be very good at extracting categories from vocalizations, but would be unable to perform family identification. Thus, a balance between the two sources of variance is necessary to perform both category generalization and family identification. We demonstrate here that our populations are grouping within-category variants (**Figure 6**) and that our population has a lower dimensionality geometry than would be expected from chance (**Figure S5**), suggesting that there is a transformation from the acoustic information either at the level of the AC or inherited from upstream regions that aids with categorical representations. A separate finding is that more single neurons are primarily specialists for either vocalization category or family ID decoding, with few neurons strongly contributing to both (**Figure 5**). Previous work has suggested that mixed-selective neurons are necessary for increasing dimensionality and that a population of purely-selective neurons limits the dimensionality of the population (Rigotti et al., 2013). It is possible that the contribution of both generalists and specialists to the decoders ensure that the AC strongly represents category, yet remains flexible enough to extract social information.

Overall, our findings provide insights into the population coding properties that support naturalistic vocalization processing. Our results are consistent with previous work showing that auditory cortex populations represent vocal categories (Mesgarani et al., 2008), specifically with a temporal code (Robotka et al., 2023), and we show that this is true even in the face of acoustic variance that conveys social identity. These category and identity representations are underpinned by a mix of both specialist and generalist neurons that work together to perform distinct computations of category generalization and family identification. Together, our findings reveal how the auditory cortex deals with naturalistic variance to provide an efficient representation of multiple types of information, permitting vocal animals to extract meaning and social identity from the same stimuli. Future investigations should build on these findings by testing how these neural computations underlie behavior decisions in naturalistic social paradigms.

## ACKNOWLEDGEMENTS

We thank Michael A. Long, Katherine I. Nagel, and J Anthony Movshon for constructive feedback and advice. We further thank the faculty of the Methods in Computational Neuroscience course at the Marine Biological Laboratories, particularly Stefano Fusi, for discussion and sharing their computational expertise. This work was supported by NIH R01DC020279 (D.H.S.) and an NSF GRFP 23-A0-00-1015499 (E.Z.).

## STAR METHODS

### EXPERIMENTAL MODEL AND SUBJECT DETAILS

Five adult Mongolian gerbils (*Meriones unguiculatus*, 2 females, 3 males) were used in the study. Gerbils were bred in an onsite vivarium, weaned at postnatal (P) day 30, and housed on a 12 hour light/dark cycle with full access to food and water. All experimental protocols were approved by New York University’s Animal Use and Welfare Committee.

### METHOD DETAILS

#### Surgeries

Animals were anesthetized with isoflurane/oxygen, secured in a stereotaxic device (Kopf), and a 4 shank 64-channel silicon probe (Cambridge Neurotech P1-236 array) was implanted in the left core auditory cortex (AC). The anterior-most shank of the probe was implanted 3.5mm anterior to lambda, 4.6mm lateral, and 1-1.2mm ventral to the surface at a 25 degree angle in the mediolateral plane. The probe was attached to a custom microdrive to allow for post-surgery advancement of the probe through AC. A ground wire was inserted into the contralateral hemisphere approximately 2mm medial and 1mm anterior from lambda.

#### Stimulus set

Vocalizations recorded from 5 families of gerbils (pups P25-27) from 7pm-7am. Parents and all pups (8 +/- 1) were placed in an enlarged cage with Alpha-Dri bedding, food, water, a roofless-house, and some bedding from the home cage. Video recordings were obtained under red LED light with a FLIR Blackfly-S camera (30 frames per second) and vocalizations were recorded at 125kHz sampling rate from 4 Avisoft microphones (placed at the four roof corners of the cage, angled towards the middle).

The signals from the 4 microphones were averaged to improve the signal-to-noise ratio (Peterson et al., 2024). We trained a network to automatically extract vocalizations from the averaged microphone signal using Deep Audio Segmenter (DAS; (Steinfath et al., 2021). To train the network, we extracted sample bouts from 8 gerbil families recorded overnight as described above. This was to ensure that the training set was not biased by the vocalizations from one particular family or by subtle changes in the background microphone signal across recording days. The network was trained by labeling bouts with one of 10 labels: alarm, noisy alarm, upward frequency modulated (ufm), noisy ufm, low-frequency, noisy-low frequency, warble, noisy-warble, chirp, and noise. This ensured that the network extracted vocalizations that were not overlapping with other vocalizations or with noise such as scratching, digging, etc. This network performed between 30-80% performance on correctly labeling variants. A second network was trained to detect noise using the noise-labeled variants from the first network, as well as the original training set. This network performed at 60% or higher for all categories and produced much fewer noise-contaminated variants (**Fig S1A-B**). The vocalization category labels used for DAS to extract variants were not used for any subsequent analyses, only for extracting vocalizations. Therefore, the model’s performance was evaluated after grouping all vocalizations, overlapping vocalizations, and noise. As demonstrated in **Fig S1B**, 80% of user-labeled vocalizations were extracted with the network. We evaluated the model’s performance for noise, vocalizations, and overlapping vocalizations using the python sklearn f1_score function (average=None).

Using the onsets and offsets output by DAS, we extracted variants from the 8 families we recorded overnight vocalizations from. The onsets and offsets from DAS showed some jitter in cropping the first or last few milliseconds of the vocalizations, so all variants were padded with 40ms to ensure that the whole vocalization was extracted. We generated spectrograms (nperseg=512, noverlap=256, spec_min_val=-8, spec_max_val = -5) from these onsets and offsets and trained a variational autoencoder to represent each spectrogram with 32 latent values (Goffinet et al., 2021). We fit a Gaussian mixture model (cov=’diag’) to cluster the vocalization latents into 200 clusters. These clusters were then manually labeled by an experimenter to be flat, stacked, spectral, flat, sweeping, short, mixed, ufm, warble, loud, and noisy. Then, we used the manual labels to pull all the GMM clusters that belonged to the broad vocal categories: Combos, DFMs, UFMs, and Warbles. For all experiments, the same trained VAE and GMM models were used on all vocalizations, so the cluster labels remained constant across all families recorded from.

The vocalizations from 4 broad categories were extracted: upward frequency modulated (UFM), warbles, downward frequency modulated (DFM), and combination vocalizations (combos). These variants were manually sorted through to check if there was at least a 100ms quiet period preceding the vocalization, to allow for a reasonable baseline period before the vocalization was presented. Variants belonging to these 4 vocalization categories were filtered with a custom python script using librosa’s soft mask vocal background subtraction. All vocalizations were normalized to a root mean squared (RMS) value of 0.004. A custom script was used to present the vocalizations with a mean inter-stimulus-interval (ISI) of .47s. This large file was divided into multiple 5 minute blocks for presentation during the experiment.

Non-vocal control stimuli were generated to test the spectrotemporal tuning properties of recorded cells. Pure tones with frequencies from 1.5-55kHz (30ms duration, 2ms rise/fall ramp) were presented randomly in 2 min blocks of 5 trials each with mean ISI of 640ms (+/- 300ms). Amplitude-modulated (AM) noise stimuli were also presented to test the rate tuning of neurons. White noise was bandpassed at 1-10kHz, 1-25kHz, 5-25kHz, 25-40kHz, or 30-55kHz and modulated at 100% depth with rates of 2, 4, 8, 16, 32, and 64Hz. AM stimuli were presented 10 times each. All stimuli were normalized to a root mean squared level of 0.004.

#### Stimulus presentation and data collection

Stimuli were presented as wavfiles using a custom script using the *pyaudio* package on a Raspberry Pi through a HiFi Berry DAC2 Pro sound card. Stimuli were amplified using a TDT z-bus stereo amplifier and presented through a Fountek NeoCd1.0 1.5" Ribbon Tweeter. At an RMS value of 0.004, the stimuli were presented at 67-75dB, as measured from either the closest position in the arena (directly below) or the furthest corner. Four ultrasonic microphones (Avisoft CM16/CMPA48AAF-5V) were synchronously recorded using a National Instruments multifunction data acquisition device (PCI-6143) via BNC connection with a National Instruments terminal block (BNC-2110). The recording was controlled with custom python scripts using the NI-DAQmx library (https://github.com/ni/nidaqmx-python) which wrote samples to disk at a 125 kHz sampling rate. The electrophysiological data was simultaneously recorded at a 12.5kHz sampling rate using a wireless data logger (White Matter LLC) that received a synchronization signal from the NIDAQ. The neural data were upsampled to 25kHz during downloading.

Implanted animals were placed in the experimental rig (22×14×14in.). The speaker was located on the roof, so sounds were presented from overhead. After 5 minutes of silence to habituate the animal, the stimulus presentation began. The order of stimulus presentation was the same every session within a block, but the block order was semi randomized across sessions. The full experimental session lasted 2-3 hours. *Electrophysiological recording and preprocessing*

Raw electrophysiological data were preprocessed using custom python scripts that bandpassed the signal from 0.2-4kHz and referenced the signal by common median subtraction. Individual action potentials from single neurons were extracted using Kilosort 2.5 and manually reviewed using Phy2. Inclusion criteria for a single unit included having at least a whole session firing rate greater than .1Hz, a contamination percentage below 20, and visually inspecting the principal components of the waveform and interspike-interval histograms for noise.

#### Acoustic analyses

To determine if the acoustics of the vocalizations could be classified based on their vocalization categories and by the identity of their family-of-origin, we used a random forest classifier. The latent values from the variational autoencoder that explained the most variance were used as input and the labels were either the vocalization categories or the family IDs from which they were recorded. The vocalization category classifiers were repeated 100 times, using either 2, 3, or 5 families’ vocalizations as input. Each iteration randomly sampled 50 syllables of each category from each family and split the variants into an 80/20 train/test set.

As described in Peterson et al. 2024, we used the *similarity_features* function from the *VocalPy* package (Nicholson, 2024), a python implementation of the Sound Analysis Pro Sound Analysis Tools library, to compute median pitch (fundamental frequency), amplitude, entropy, frequency modulation, and goodness of pitch. In addition, spectral flatness was computed using the python *librosa* package and duration was computed from the audio. Start and stop frequencies were computed by taking the median fundamental frequency within the first and last thirds of each vocalization.

Bandwidth was computed by taking the difference between the start and stop frequencies.

#### Electrophysiological analyses

##### Quantification of responsiveness to vocal categories, family identities, tones, and noise

Firing rates were computed by summing the number of spikes in a given epoch window and dividing by the length of this window. Epochs for which firing rate was computed included the spontaneous period (100ms before the vocalization’s background started), the pre-vocalization background period (i.e. pad), and the duration of the vocalization itself. A neuron was counted as responsive to a vocal category if the firing rate during the vocalization epoch across all trials of that vocalization was significantly different from the firing rates of both the spontaneous epoch and the pad epoch (paired t-test with Bonferroni correction; *p*<2.5e-3). From these analyses, we counted the number of cells that were significantly responsive to 0, 1, 2, 3, or all 4 categories, as well as which categories these neurons were responsive to.

To determine whether more neurons were responsive to an animal’s own family than the other families, we quantified the responsiveness of cells specifically to each family ID. If a neuron was responsive to a given category, then we compared the spontaneous, pad, and evoked firing rates and the for each family of that category individually. If the evoked firing rates were significantly different from both the spontaneous and pad (paired t-test with Bonferroni correction; *p*<3.3e-3), the unit was counted as ‘family responsive’.

We also computed responsiveness to tones and noise. For tones, the firing rates 100ms before tone-onset were compared to the firing rates from tone onset to 50ms after tone offset (to account for some neurons displaying prolonged tone firing). A neuron was counted as tone-responsive if there was at least one tone frequency for which the evoked firing rate was significantly different than the spontaneous firing rate (paired t-test with Bonferroni correction; *p*<2.6e-4). The neuron’s best frequency was determined for tone-responsive units as the frequency that elicited the maximum change in firing compared to baseline. Noise-responsiveness was determined by comparing the 100ms period before noise onset and the firing rate during the full 1 second amplitude-modulated noise stimulus (paired t-test with Bonferroni correction; *p*<3.3e-4).

##### Selectivity index

For each of the vocal-responsive units, a selectivity index (as described in Sharma et al., 2024) was computed to determine how specific a neuron’s response was for 1 versus all categories. An SI=1 indicates that a neuron only responds to 1 category and not to any of the others, while SI=0 indicates equal firing to all four categories. The formula is the following:

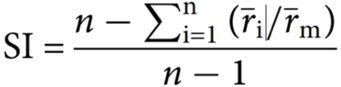

where *n* represents the number of categories (4) or families (3), r^i^ represents the absolute value of the difference between the mean category/family firing rate and the mean spontaneous category/family firing rate, and r_m_ is the maximum of r_i_ for all i. For vocalization category selectivity (**Fig 2G**), the mean category firing rates were used, pooling across family ID). For family ID selectivity, the mean family firing rates within a given category were used. Note that only category-responsive neurons were used to compute family ID selectivity for a given category. To determine if a unit was family-modulated, we used a one-way ANOVA to test if the firing rates were significantly different across families (one-way ANOVA, *p*<2.5e-3). The number of units that were responsive to family identity for at least 1 category was divided by the total number of vocal-responsive units to determine the percentage of the responsive population that was family-modulated.

##### Population decoders

Classification of single trial, z-scored 30ms-binned PSTHs was performed using a linear support vector machine (SVM) using the Python *sklearn* package, with parameters C=1.0, tol=1e-3, penalty=’l1’, dual=False, and max_iter=10000. The shape of the decoder input had the rows, or features, N_neurons_xB_timebins_, and columns, or samples, T_trials_. Since one family was always the subject animal’s own family, only 2 families were used for identity decoding performance in this analysis (so that we could include all variants that were presented to all neurons in the dataset). The decoder was performed 100 times, sampling 300 trials from each category and family per iteration to ensure equal sampling of the dataset. On a given iteration, the data were separated 80/20 into train and test sets, and the percent correct of the decoder on the test set was saved. To ensure fair comparison of vocalization category and family ID decoding, the same sampled trials that were used for decoding family ID were further subsampled so that there were always 300 trials per class (for family: 300 from Fam1 and 300 from Fam2, for vocalization category: 150 from each family, so that there were 300 trials per class). The exact same decoding analysis was performed using either a pseudopopulation of all units, a pseudopopulation of all vocal-responsive units, or using each individual gerbil’s vocal-responsive units. We also performed the decoding analysis as described using the z-scored firing rates rather than the z-scored PSTHs (input: N_neurons_xT_trials_).

We determined the effect of population size on decoding performance by performing the decoding as described, but also subsampling increasing numbers of neurons in the population. We sampled 2, 5, 10, 25, 50, 100, 200, and 316 neurons 20 times each. The mean performance was fit with a sigmoidal function by bootstrapping the individual iteration performances 500 times. The sigmoid function was:Where x is the number of neurons, y is the SVM performance, L is the range of the function, k is the steepness of the curve, x_0_ is the inflection point, and b is the lower asymptote of the function. The bootstrap was initialized with x_0_ as the number of neurons needed for the average performance across all neuron samplings and b was constrained with a lower bound of chance performance (25% for vocalization category, 50% for family ID).

We further analyzed the weights of neurons in the PSTH SVM to determine the relative contribution of individual neurons to either category or family ID decoding. We performed the SVM 1000 times, as well as a null decoder that was trained on shuffled trial labels. and saved the coef_ for each iteration. Since the PSTH SVM input was N_neurons_xB_bins_, each iteration produced 8 weights per neuron. Further, we were interested in the relative contribution of each cell, so we computed the mean absolute value of these weights for each cell in a given iteration. We then divided the mean(abs(weight)) of each cell by the sum of all the weights to determine the percent contributed by each neuron. A neuron was determined to significantly contribute to a decoder if its ‘percent contribution’ across the 1000 iterations was significantly higher than in the null decoder (paired t-test with Bonferroni correction; *p*<2.5e-3).

##### Categorization analysis: acoustics versus neural performance

To determine if AC populations displayed a boundary effect for the representation of vocalization categories, we directly compared the performance of the SVM on classifying the acoustics versus the acoustically-evoked neural activity. For each vocalization category, we first used the latent values to sample 2 variants belonging either to the same (‘within-category’) or a different vocalization category (‘across-boundary’). Next, the cosine similarity between all the variants of that vocalization category and the 2 sampled variants were calculated to select the 24 variants most acoustically similar to the sampled ones. Then, an SVM was trained on either the z-scored acoustic features of those 50 variants or the trial-averaged z-scored 30ms PSTHs. This procedure was repeated 500 times for each vocalization category, with 5 cross-validation shuffles of the train/test set for each 50-variant sample.

#### Histological electrode site confirmation

The experimental endpoint was determined either by the microdrive reaching its deepest point in the tissue such that no new AC neurons could be recorded from, or the recording SNR dropping to a level where single units could no longer be sorted. Animals were deeply anesthetized with isoflurane and overdosed with 0.3mL euthasol. Once the animal stopped breathing but before the heart stopped beating, a transcardial perfusion was performed with 10mL of saline and 20mL of 10% formalin. The brain was extracted and stored in 10% formalin for at least 2 days. The brain was notched on the right hemisphere and placed in agar so that it could be sectioned into 60um sections using a vibratome (Leica). Sections were mounted onto gelatin-subbed slides and left to dry for 24 hours. Sections were then stained with 0.25% thionin and coverslipped with Permount. After drying, sections were imaged to locate the probe. Probe sites were aligned to the Ratdke-Schuller gerbil stereotaxic atlas (Radtke-Schuller et al., 2016) to determine the exact implantation sites.

